# Statistical Analysis of Variability in TnSeq Data Across Conditions Using Zero-Inflated Negative Binomial Regression

**DOI:** 10.1101/590281

**Authors:** Siddharth Subramaniyam, Anisha Zaveri, Michael A. DeJesus, Clare Smith, Richard E. Baker, Sabine Ehrt, Dirk Schnappinger, Christopher M. Sassetti, Thomas R. Ioerger

## Abstract

Deep sequencing of transposon mutant libraries (or TnSeq) is a powerful method for probing essentiality of genomic loci under different environmental conditions. Various analytical methods have been described for identifying conditionally essential genes whose tolerance for insertions varies between two conditions. However, for large-scale experiments involving many conditions, a method is needed for identifying genes that exhibit significant variability in insertions across multiple conditions. In this paper, we introduce a novel statistical method for identifying genes with significant variability of insertion counts across multiple conditions based on Zero-Inflated Negative Binomial (ZINB) regression. Using likelihood ratio tests, we show that the ZINB fits TnSeq data better than either ANOVA or a Negative Binomial (in a generalized linear model). We use ZINB regression to identify genes required for infection of *M. tuberculosis* H37Rv in C57BL/6 mice. We also use ZINB to perform a retrospective analysis of genes conditionally essential in H37Rv cultures exposed to multiple antibiotics. Our results show that, not only does ZINB generally identify most of the genes found by pairwise resampling (and vastly out-performs ANOVA), but it also identifies additional genes where variability is detectable only when the magnitudes of insertion counts are treated separately from local differences in saturation, as in the ZINB model.

## 1 Background

Deep sequencing of transposon mutant libraries (or TnSeq) is a powerful method for probing the essentiality of genomic loci under different environmental conditions [1]. In a transposon (Tn) mutant library made with a transposon in the *mariner* family, like Himar1, insertions generally occur at approximately random locations throughout the genome, restricted to TA dinucleotides [2]. The absence of insertions in a locus is used to infer conditional essentiality, reflecting depletion of those clones from the population due to inability to survive the loss of function in such conditions. If loss of function leads to a significant growth impairment, these genes are typically referred to as ‘growth-defect’ genes instead. While the abundance of clones with insertions at different sites can be profiled efficiently through deep sequencing [3], there are a number of sources of noise that induce a high degree of variability in insertion counts at each site, including: variations in mutant abundance during library construction [4], stochastic differences among replicates [5], biases due to sample preparation protocol and sequencing technology [6], and other effects. Previous statistical methods have been developed for quantitative assessment of essential genes in single conditions, as well as pairwise comparisons of conditional essentiality. Statistical methods for characterizing essential regions in a genome include those based on tests of sums of insertion counts in genes [7], gaps [8], bimodality of empirical distributions [9], non-parametric tests of counts [10], Poisson distributions [11], and Hidden Markov Models [12, 13]. Statistical methods for evaluating *conditional* essentiality between two conditions include: estimation of fitness differences [14], permutation tests on distribution of counts at individual TA sites (resampling in TRANSIT [15]), Mann-Whitney U-test [16], and linear modeling of condition-specific effects (i.e. log-fold-changes in insertion counts) at individual sites, followed by combining site-level confidence distributions on the parameters into gene-level confidence distributions (TnseqDiff [17]).

Recently, more complex TnSeq experiments are being conducted involving larger collections of conditions (such as assessment of a library under multiple nutrient sources, exposure to different stresses like a panel of antibiotics, or passaging through multiple animal models with different genetic backgrounds) [18, 19, 20, 21]. Yang et al. [22] has also looked at temporal patterns of changes in insertion counts over a time-course. A fundamental question in such large-scale experiments is to determine which genes exhibit statistically significant variability across the panel of conditions. A candidate approach might be to perform an ANOVA analysis of the insertion counts to determine whether there is a condition-dependent effect on the means. However, ANOVA analyses rely on the assumption of normality [23], and Tn insertion counts are clearly not Normally distributed. First, read-counts are non-negative integers; second, there are frequently sporadic sites with high counts that influence the means; third, most Tn libraries are sub-saturated, with a high fraction of TA sites not being represented, even in non-essential regions. This creates an excess of zeros in the data (sites were no insertion was observed), and this makes it ambiguous whether sites with a count of 0 are biologically essential (i.e. depleted during growth/selection) or simply missing from the library. Monte Carlo simulations show that applying ANOVA to data with non-normally distributed residuals can result in an increased risk of type I or type II errors, depending on degree and type of non-normality [23]. An alternative method for assessing variability might be to use a non-parametric test of the differences between means by permuting the counts and generating a null distribution (as in the “resampling test” in TRANSIT [15]). However, this is limited to pairwise comparisons, and attempting to run resampling for all pairwise comparisons between conditions to identify genes that show some variation does not scale up well as the number of conditions grows.

In this paper, we introduce a new statistical method for identifying genes with significant variability of insertion counts across multiple conditions based on Zero-Inflated Negative Binomial (ZINB) regression. The ZINB distribution is a mixture model of a Negative Binomial distribution (for the magnitudes of insertion counts at sites with insertions) combined with a “zero” component (for representing the proportion of sites without insertions). ZINB regression fits a model for each gene that can be used to test whether there is a condition-dependent effect on the magnitudes of insertion counts or on the local level of saturation in each gene. Separating these factors increases the statistical power that ZINB regression has over resampling for identifying varying genes (since resampling just tests the differences in the means between conditions - zeros included). Importantly, our model includes terms to accommodate differences in saturation among the datasets to prevent detecting false positives due to differences between libraries.

Another advantage of the ZINB regression framework is that it allows incorporation of additional factors as covariates in analyzing variability across multiple conditions, to account for effects dependent on relationships among the conditions, such as similar treatments, time-points, host genotypes, etc.

Using several TnSeq datasets from *M. tuberculosis* H37Rv, we show that, in pairwise tests (between two conditions), the genes detected by ZINB regression are typically a superset of those detected by resampling and hence is more sensitive. More importantly, ZINB regression can be used to identify varying genes across multiple (≥ 3) conditions, which contains most of the genes identified by pairwise resampling between all pairs (and is more convenient and scalable). Furthermore, ZINB regression vastly out-performs ANOVA, which often identifies only around half as many genes with significant variability in insertion counts.

## 2 Methods

### 2.1 ZINB Model

Essential genes are likely to have no insertions or very few counts (because mutants with transposon insertions in those regions are not viable), while non-essential genes are likely to have counts near the global average for the dataset. Insertion counts at TA sites in non-essential regions are typically expected to approximate a Poisson distribution. This expectation is based on a null model in which the expected fraction of insertions at a site is determined by the relative abundance of those clones in the library, and the observed counts in a sequencing experiment come from a stochastic sampling process. This process is expected to follow a multinomial distribution [24], which is approximated by the Poisson for sufficiently large numbers of reads (total dataset size) [25].

Let *Y* = {*y_g,c,i,j_*} represent the set of observed read counts for each gene *g*, in condition *c* ∈ {*c*_1_..*c_n_*}, at TA site *i* = 1..*N_g_*, for replicate *j* = 1..*R_c_*. We are interested in modeling the gene- and condition-specific effects on the counts, *p*(*y|g, c, i, j*). We treat the observations at individual TA sites and in different replicates as independent identically-distributed (i.i.d.), samples drawn from the distribution for the gene and condition:

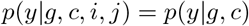

Read-count data is often modeled using the Negative Binomial (NB) distribution [25]. The NB distribution can be thought of as a Poisson distribution with over-dispersion, resulting from an extra degree of freedom:

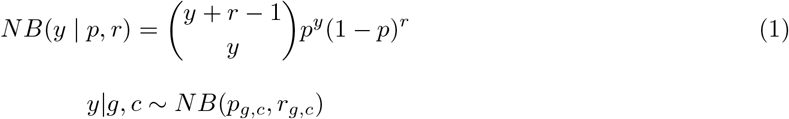

where *p* is a success probability (i.e. of a mutant getting a transposon insertion at a particular site), and *r*, often called a size parameter, represents the dispersion. Unlike the Poisson distribution, which has a single parameter λ = 1/*p*, and for which the variance is restricted to equal the mean, the extra parameter in NB allows for fitting counts with a variance greater or less than expected (i.e. different from the mean). The NB distribution converges to a Poisson as *r* → ∞ [26]. A common re-parameterization of the NB distribution is to specify the distribution based on the mean, *μ*, and the dispersion parameter, *r*, which then determines the success probability, *p*, through the following relationship:

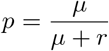

In practice, TnSeq data often has an excess of empty sites (TA sites with counts of 0), exceeding those that would be expected under a typical NB distribution. Because essential genes typically constitute only 10 — 20% of the genome in most organisms, a library with transposon insertions at 50% of its sites (i.e. 50% saturation) would mean that even non-essential genes will have a large portion of sites missing (i.e. equal to zero). Thus, while the NB distribution may be sufficient to model counts in other domains, TnSeq requires more careful consideration.

One way to solve this problem is to model the read-counts for a gene *g* and condition c as coming from a Zero-Inflated Negative Binomial distribution (ZINB) instead:

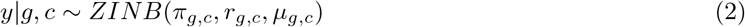

where

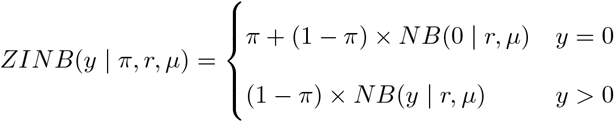

Here the *π* parameter represents the probability that a count of zero is extraneous (i.e. does not belong to the NB distribution), and can be interpreted as similar to the probability that an empty site is essential (i.e. empty due to fitness costs incurred through its disruption, rather than stochastic absences). In this way, both read-counts (through the *r* and *μ* parameters of the NB distribution) and insertion density (through *π*) can be used to differentiate genes that are essential in one condition and non-essential in another.

### 2.2 Generalized Linear Model

To capture the conditional dependence of the ZINB parameters (*μ, r, π*) on the experimental conditions, we adopt a linear regression (GLM) approach, using a log-link function. This is done independently for each gene *g*. We use *Y_g_* = {*y_g_*,.,.,.} to represent the set of all observed counts in gene *g* at any TA site, in any condition, in any replicate. The vector of expected means *μ_g_* of the ZINB distribution (non-zero component) for each observation in gene g is expressed as:

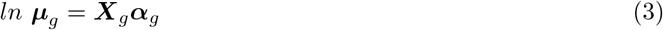

where ***X**_g_* is a binary design matrix indicating the experimental condition for each observation (sample treatments for each insertion count at each TA site in gene *g*), and ***α**_g_* is a vector of coefficients for each condition. For *m* observations and *n* conditions, the size of ***X**_g_* will be *m* × *n* and the size of ***α**_g_* will be *n* × 1. Hence, there will be *n* coefficients for each gene, one for estimating the mean non-zero count for each condition. The conditional expectations for the non-zero means for each condition can be recovered as: 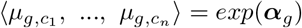.

If additional covariates distinguishing the samples are available, such as library, timepoint, or genotype, they may be conveniently incorporated in the linear model with an extra matrix of covariates, ***W**_g_* (*m × k* for *k* covariates), to which a vector of *k* parameters ***β**_g_* will be fit:

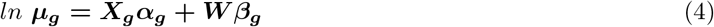

For the dispersion parameter of the NB, *τ* (or size parameter *r* = 1/*τ*), we assume that each gene could have its own dispersion, but for simplicity, we assume that it does not differ among conditions. Hence, it is fitted by a common intercept:

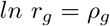

Finally, for the zero-inflated (Bernoulli) parameter, *π*, we fit a linear model depending on condition, with a logit link function (a conventional choice for incorporating probabilistic variables bounded between 0 and 1 as terms in a linear model):

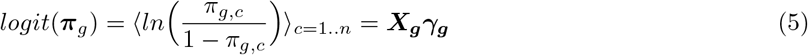

Thus each gene will have its own local estimate of insertion density in each condition, *π_g,c_* = *exp*(*γ_g,c_*)/(1 + *exp*(*γ_g,c_*)). In the case of covariates, *logit*(***π**_g_*) = ***X_g_γ_g_*** + ***W_g_δ_g_***, where ***W_g_*** are the covariates for each observation, and ***δ**_g_* are the coefficients for them.

Putting these all together:

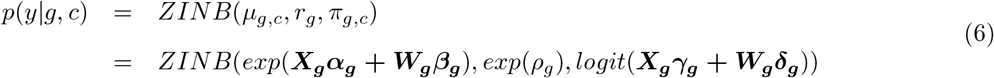

The parameters of the GLM can be solved by maximum-likelihood using iteratively re-weighted least squares (IWLS). In this work, we use the *pscl* package in R [27].

### 2.3 Correcting for Saturation Differences Among TnSeq Datasets

An important aspect of comparative analysis of TnSeq data is the normalization of datasets. Typically, read-counts are normalized such that the total number of reads is balanced across the datasets being compared. Assuming read-counts are distributed as a mixture of a Bernoulli distribution (responsible for zeros) and another distribution, *g*(*x*), responsible for non-zero counts i.e.,

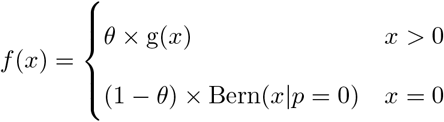

then the expected value of this theoretical read-count distribution (with mixture coefficient *θ*) is given by:

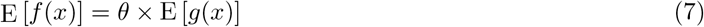

The expected value of such a distribution can be normalized to match that of a another dataset, *f_r_*(*x*), (such as reference condition, with saturation *θ_r_*) by multiplying it by a factor, *w*, defined in the following way:

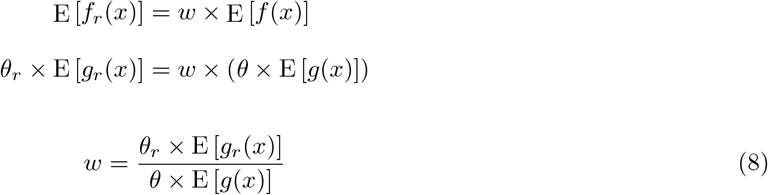

This guarantees that the expected value in read-counts is the same across all datasets. TTR normalization (i.e. total trimmed read count, the default in TRANSIT [15]) estimates **E**[*g*(*x*)] in a robust manner (excluding the top 1% of sites with highest counts, to reduce the influence of outliers, which can affect normalization and lead to false positives).

While TTR works well for methods like resampling (which only depend on the expected counts being equivalent under the null-hypothesis), it does not work well for methods designed to simultaneously detect differences in both the local magnitudes of counts (non-zero mean) and the saturation (fraction of non-zero sites) such as ZINB. This is because TTR in effect inflates the counts at non-zero sites in datasets with low saturation, in order to compensate for the additional zeros (to make their expected values equivalent). This would cause genes to appear to have differences in (non-zero) mean count (*μ_g,a_* vs *μ_g,b_*), while also appearing to be less saturated (*π_g,a_* vs *π_g,b_*), resulting in false positives.

To correct for differences in saturation, we incorporate *offsets* in the linear model as follows. First, assume there are *d* datasets (combining all replicates over all conditions). Let the statistics of each dataset be represented by a *d* × 1 vector of non-zero means, **M** (genome-wide averages of insertion counts at non-zero sites), and a *d* × 1 vector of the fraction of sites with zeros in each each dataset, ***Z***. For the m observations (insertion counts at TA sites) in gene *g*, let *D_g_* be the binary design matrix of size *m × d* indicating the dataset for each observation. Then the linear equations above can be modified to incorporate these offsets (a specific offset for each observation depending on which dataset it comes from).

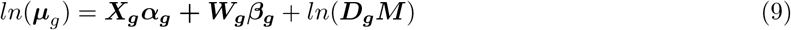

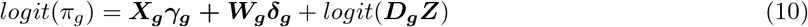

Note that ***M*** and ***Z*** are just vectors of empirical constants in the linear equation, not parameters to be fit. Hence the fitted coefficients (***α_g_, β_g_,γ_g_, δ_g_***) are effectively estimating the *deviations* in the local insertion counts in a gene relative to the global mean and saturation for each dataset. For example, if observation *X_g,c,i,j_* comes from dataset *d* (where *i* and *j* are indexes of TA site and replicate), and the global non-zero mean of that dataset is *M_d_*, then *exp*(*X_g_α_g_*) estimates the ratio of the expected mean insertion count for gene *g* in condition *c* to the global mean for dataset *d* (ignoring covariates):

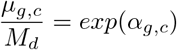

### 2.4 Statistical Significance

Once the ZINB model is fit to the counts for a gene, it is necessary to evaluate the significance of the fit. T-tests could be used to evaluate the significance of individual coefficients (i.e. whether they are significantly different from 0). However, for assessing whether there is an overall effect as a function of condition, we compare the fit of the data *Y_g_* (a set of observed counts for gene *g*) to a simpler model -ZINB without conditional dependence - and compute the difference of log-likelihoods (or log-likelihood ratio):

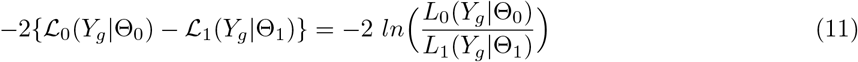

where the two models are given by:

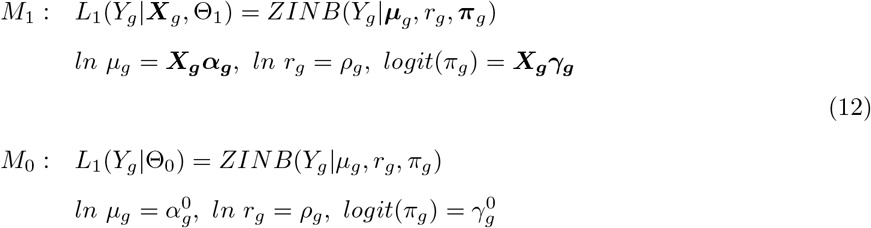

where 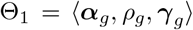 and 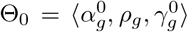 are the collections of parameters for the two models, and where 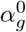 and 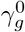 in *M*_0_ are just scalars fitted to the grand mean and saturation of the gene over all conditions. The likelihood ratio statistic above is expected to be distributed as *χ*^2^ with degrees of freedom equal to the difference in the number of parameters (Wilks’ Theorem):

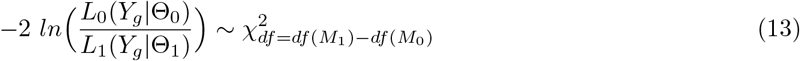

For the condition-dependent ZINB model (*M*_1_), the number of parameters is 2*n* + 1 (for length of ***α**_g_* and *γ_g_* plus *ρ_g_*). For the condition-independent ZINB model (*M*_0_), there are only 3 scalar parameters 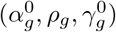 used to model the counts pooled across all conditions. Hence *df* = 2*n* +1 – 3 = 2(*n* – 1). The point of the test is to determine whether the additional parameters, which should naturally improve the fit to the data, are justified by the extent of increase in the likelihood of the fit. The cumulative of the *χ*^2^ distribution is used to calculate p-values from the log-likelihood ratio, which are then adjusted by the Benjamini-Hochberg procedure [28] to correct for multiple tests (to limit the false-discovery rate to 5% over all genes in the genome being tested in parallel).

Importantly, if a gene is detected to be conditionally-essential (or have a conditional growth defect), it could due to either a difference in the mean counts (at non-zero sites), or saturation, or both. Thus the ZINB regression method is capable of detecting genes that have insertions in approximately the same fraction of sites but with a systematically lower count (e.g. reduction by X%), possibly reflecting a fitness defect. Similarly, genes where most sites become depleted (exhibiting reduced saturation) but where the mean at the remaining sites (perhaps at the termini) remains about the same would also be detectable as conditional-essentials.

### 2.5 Covariates and Interactions

If the data include additional covariates, then the *W* terms will be included in the regressions for *both* models *M*_1_ and *M*_o_:

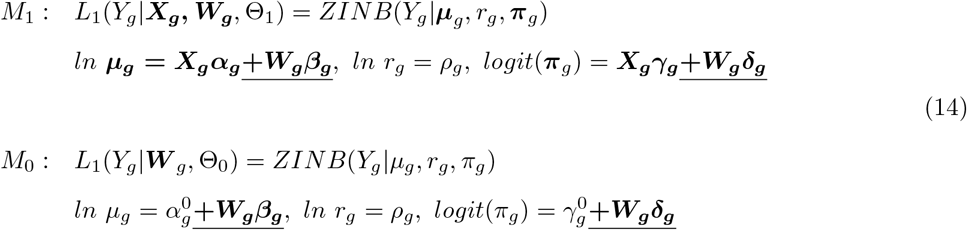

In this way, the covariates *W* will increase the likelihoods of both models similarly, and the LRT will be evaluating only the improvement of the fits due to the conditions of interest, *X*, i.e. the residual variance explained by *X* after taking known factors *W* into account. Although the number of parameters in both models will increase, the difference in degrees of freedom will remain the same.

If the covariates represent attributes of the samples that could be considered to interact with the main condition, then one can test for *interactions* by including an additional term in the regression. For example, suppose the main distinction of interest among the samples is strain (e.g. knockout vs wildtype), but the samples have been cultured over several times points (e.g. 1-3 weeks), then we might naturally expect that there will be variability across all 6 conditions (treated independently). Some genes might exhibit a gradual increase or decrease in counts over time. The question we really want to ask is whether the *slopes* differ significantly depending on the strain. More generally, are the differences in insertion counts among conditions significantly influenced by the value of the interacting variable?

Interactions can be incorporated in the ZINB regression model by including the *product* of the conditions with the interacting covariates in the regression for ***M***_1_.

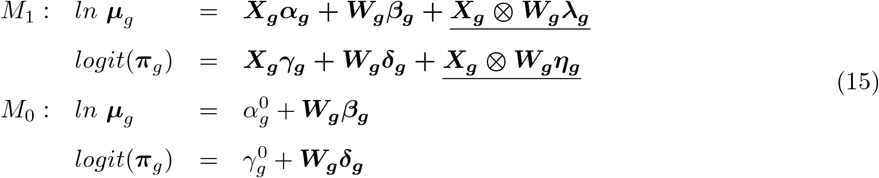

where ***X_g_*** ⊗ ***W_g_*** represents column-wise products for each pair of columns in ***X**_g_* and ***W**_g_* (resulting in a matrix of dimensions *m* × (*n* · *k*) for *n* conditions and *k* interaction variables). Thus, if there is a general trend in the counts for a gene over time, it will be captured by the coefficients of ***W**_g_* (vectors *β_g_* and ***δ**_g_*), included in both models. However, if the variables ***X**_g_* and ***W**_g_* interact, then the coefficients of the product term (**λ**_*g*_ and ***η**_g_*) will be non-zero, allowing the slopes to differ between the strains. Importantly, because the objective is to test for the significance of the interaction, in the likelihood-ratio test, the additive term for the covariate is retained in the null model but not the product, thus assessing the specific impact of the interaction on reducing the likelihood, while factoring out the information (i.e. general trend) attributable to the interaction variable on its own (independent of the main condition).

### 2.6 Availability

The methods described in this paper have been implemented in TRANSIT [15], which is publicly available on GitHub (http://saclab.tamu.edu/essentiality/transit/) and can be installed as a python package *(tnseq-transit*) using *pip*.

## 3 Results

### 3.1 Likelihood Ratio Tests for Suitability of ZINB as a Model for TnSeq Data

To establish the suitability of ZINB as a model for TnSeq data, we compared it to ANOVA and Negative Binomial (without special treatment of zeros) using likelihood ratio tests. The data we used for these tests consisted of 2 replicates of an *M. tuberculosis* H37Rv TnSeq library grown on glycerol compared to 3 replicates grown on cholesterol [29]. This data was originally used to identity genes in the H37Rv genome that are necessary to catabolize cholesterol, a unique carbon source available within the restricted intracellular environment of macrophages, on which growth and survival of the bacilli depends [30]. The data (insertion counts at TA sites) were normalized by the TTR method [15].

First, we compared ZINB regression to simple ANOVA (based on a generalized linear model using Gaussian likelihood functions). Both models were used to fit the insertion-count observations at the TA sites in each gene, conditioned on the carbon source (glycerol vs. cholesterol). ZINB had higher likelihood than ANOVA for all genes (except five, for which they were nearly equal). Because ZINB and ANOVA are not nested models, we used the Vuong test [31] to evaluate statistical significance of the difference in likelihoods. Furthermore, we applied the Benjamini-Hochberg procedure to adjust the p-values for an overall false-discovery rate (FDR) of 5%. ZINB was found to produce a significantly better fit than ANOVA for 3185 out of 3282 genes (97%, using *p_adj_* < 0.05 as a criterion).

Next, we performed a likelihood ratio test (LRT) of ZINB regression compared to regular NB (as a generalized linear model). Because ZINB has more parameters (and these are nested models), the likelihood for ZINB was again higher than NB for nearly every gene. To evaluate which differences were significant, correcting for the different number of parameters, we computed p-values of the log-likelihood ratio using the *χ*^2^ distribution, with degrees of freedom equal to the difference in number of model parameters (*df* = 5 — 3 = 2). After FDR-correction, ZINB fit the data significantly better than NB for 2796 genes out of 3282 (85%) genes evaluated. For the rest of the genes, the likelihoods of the two models were indistinguishable. This supports the hypothesis that modeling the fraction of sites with no insertions (“zeros”) separately from the magnitudes of counts at sites with insertions enables ZINB to fit TnSeq data better.

### 3.2 Pairwise Comparisons of Conditional Essentiality Using ZINB

We evaluated ZINB, resampling, and ANOVA on data from an *M. tuberculosis* TnSeq library grown in-vitro compared to infections in a mouse model. A high-saturation Himar1 Tn library generated in H37Rv was inoculated into six C57BL/6 mice (8-12 week old males, obtained from Jackson Laboratory, Bar Harbor, ME) via the intravenous route at a dose that deposits a representative sample of the library (>100,000 CFU) in the spleen. After four weeks, the bacteria present in the spleen of each animal were recovered by plating on 7H10 agar (with kanamycin). As a control, the original library was replated in parallel. A total of 0.4-1.5 million reads was mapped to TA sites for each sample, and all samples had ~ 50% saturation (all but one were in the 42-58% range; see Table 1). The data was normalized using TTR (Trimmed Total Read-count) normalization [15], and the mean count of all datasets after normalization was uniform, around 100.

**Table 1:**
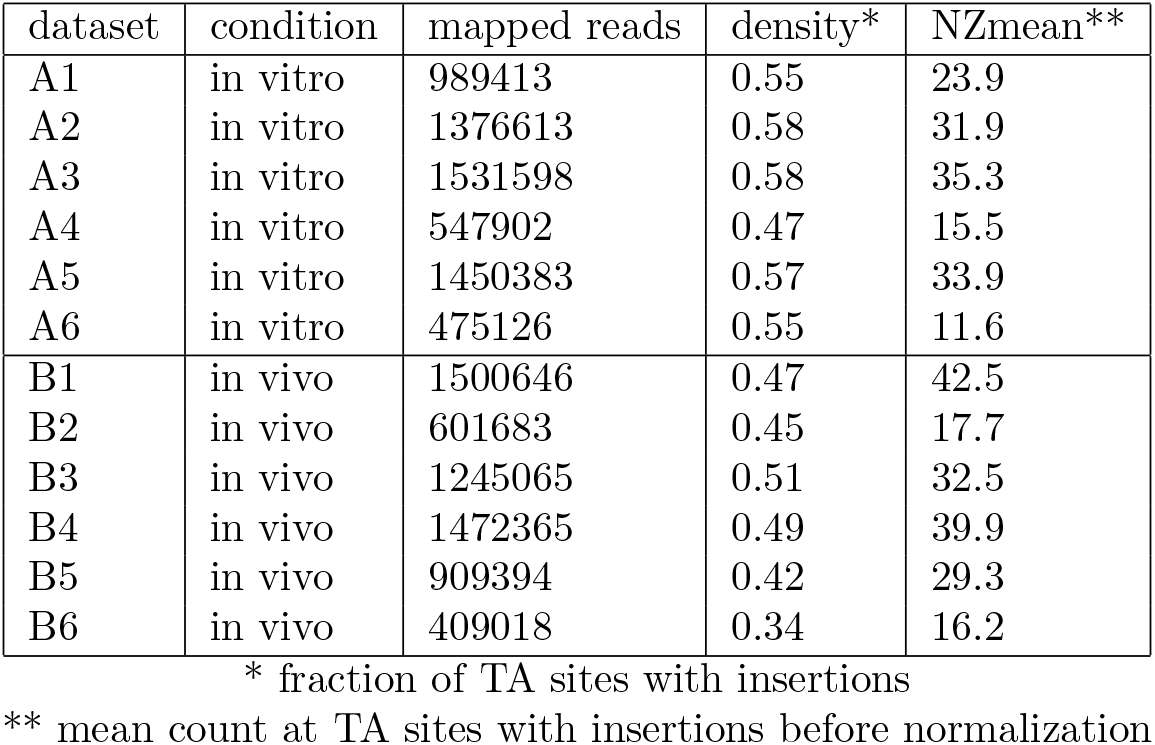
Statistics of TnSeq datasets

| dataset | condition | mapped reads | density* | NZmean** |
| --- | --- | --- | --- | --- |
| A1 | in vitro | 989413 | 0.55 | 23.9 |
| A2 | in vitro | 1376613 | 0.58 | 31.9 |
| A3 | in vitro | 1531598 | 0.58 | 35.3 |
| A4 | in vitro | 547902 | 0.47 | 15.5 |
| A5 | in vitro | 1450383 | 0.57 | 33.9 |
| A6 | in vitro | 475126 | 0.55 | 11.6 |
| B1 | in vivo | 1500646 | 0.47 | 42.5 |
| B2 | in vivo | 601683 | 0.45 | 17.7 |
| B3 | in vivo | 1245065 | 0.51 | 32.5 |
| B4 | in vivo | 1472365 | 0.49 | 39.9 |
| B5 | in vivo | 909394 | 0.42 | 29.3 |
| B6 | in vivo | 409018 | 0.34 | 16.2 |

\* fraction of TA sites with insertions
\*\* mean count at TA sites with insertions before normalization

When ZINB regression method was run on the two conditions (in vitro vs. in mice), 237 conditional essentials were identified. This included genes well-known to be essential *in vivo* [32], including the Mce4 cluster, biotin biosynthesis *(bioABDF1),* ESX-1, the NRPS (non-ribosomal peptide synthase) cluster (Rv0096-Rv0101), and cholesterol catabolism genes (e.g. FadE5, *bpoC, hsaD).* Some genes involved in mycobactin-dependent iron acquisition *(irtAB, mmpL4/S4*) were essential in vivo, though none of the 14 subunits of mycobactin synthase (Mbt) were. A possible explanation is that mutants with disruptions in Mbt genes are importing extracellular mycobactin produced by other mutants at the site of infection with insertions in genes other than Mbt synthase. In contrast to infections with a homogeneous knockout mutant of genes like *MbtD*, mycobactin synthase transposon mutants in the Tn library can survive in vivo because it is a heterogeneous pool. However, individual clones with defects in mycobactin secretion/uptake (e.g. Tn insertions in *irtAB* and *mmpL4/S4*) cannot survive, despite the availablility of mycobactin in the environment.

The results of ZINB can be compared to the permutation test (‘resampling’ in TRANSIT), which is a non-parameteric comparison of the difference in mean counts for each gene between the two conditions. Resampling yielded 186 genes with significant differences between in-vitro and in-vivo. (P-values for all tests were corrected for a false-discovery rate of < 5% using the Benjamini-Hochberg procedure [28]). Almost all of these (160, 86%) were contained in the hits from ZINB (see Figure 1). Only 26 genes identified by resampling were not detected by ZINB. Many of these were marginal cases; 21 of 26 had ZINB adjusted p-values between 0.05 and 0.2.

**Figure 1:**
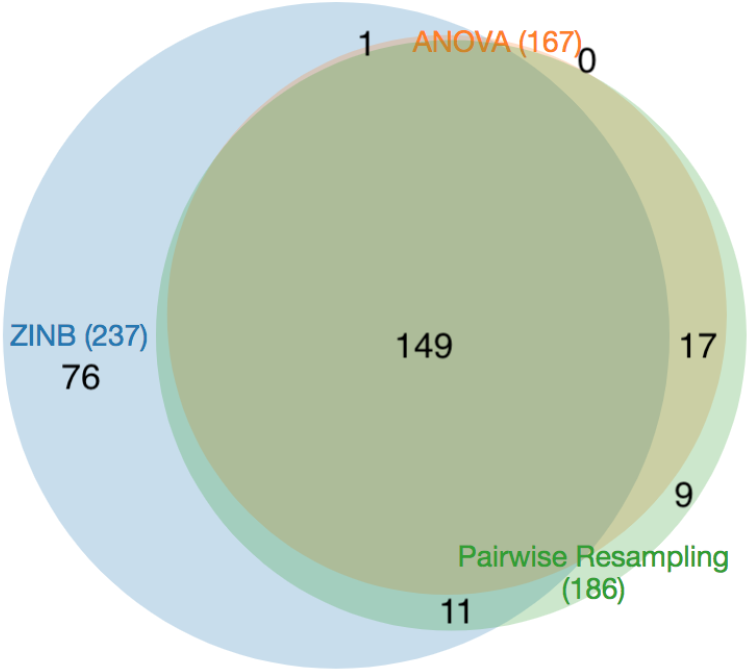
Venn diagram of conditional essentials (*qval* < 0.05) for three different methods: resampling, ANOVA, and ZINB.

ANOVA was also applied to the same data, and it only detected 167 genes with significant variability between the two conditions. The genes detected by ANOVA were almost entirely contained in the set of genes detected by resampling (166 out of 167), but resampling found 20 more varying genes. In comparison, ANOVA only finds 63% of the varying genes detected by ZINB (150 out of 237). We speculate that the lower sensitivity of ANOVA is due to the non-normality of insertion-count data, which is supported by simulation studies [23], whereas resampling, being a non-parametric test, does not require normality.

The advantage of ZINB is that it is capable of detecting more conditional essentials because it can take into account changes in either the local magnitude of counts or local insertion density. It detects 76 more conditional essentials and growth-defect genes than resampling, and 88 more than ANOVA. Among these are genes in the Mce1 cluster (specifically *mce1B, mce1C,* and *mce1F*, see Figure 2). Mce1 (Mammalian Cell Entry 1) is a membrane transporter complex that has been shown to be essential for growth in vivo (e.g. knockout mutants are attenuated for survival in mice [32, 33]). The Mce1 locus spans Rv0166-Rv0178 (as an operon), containing *mce1A-mce1F*, which are 5 subunits that form a membrane complex [34]; the rest of the proteins in the locus (*yrb1AB, mam1ABCD)* are also membrane-associated [35]. The Mce1 genes show a modest reduction in counts (~ 25% reduction; mean log_2_-fold-change=-0.2, range=-0.87..0.21), which was not sufficient to meet the adjusted p-value cutoff for resampling. However, the genes also exhibit a noticable reduction in local saturation in this locus (from ~ 88% saturation in-vitro to ~ 61% in-vivo on average), and the combination of these two depletion effects is sufficient to make them significant in the ZINB model. This is consistent with our understanding of the biological role of Mce1, which acts as a transporter to enhance uptake of fatty acids as a carbon source from the host environment [36, 37].

**Figure 2:**
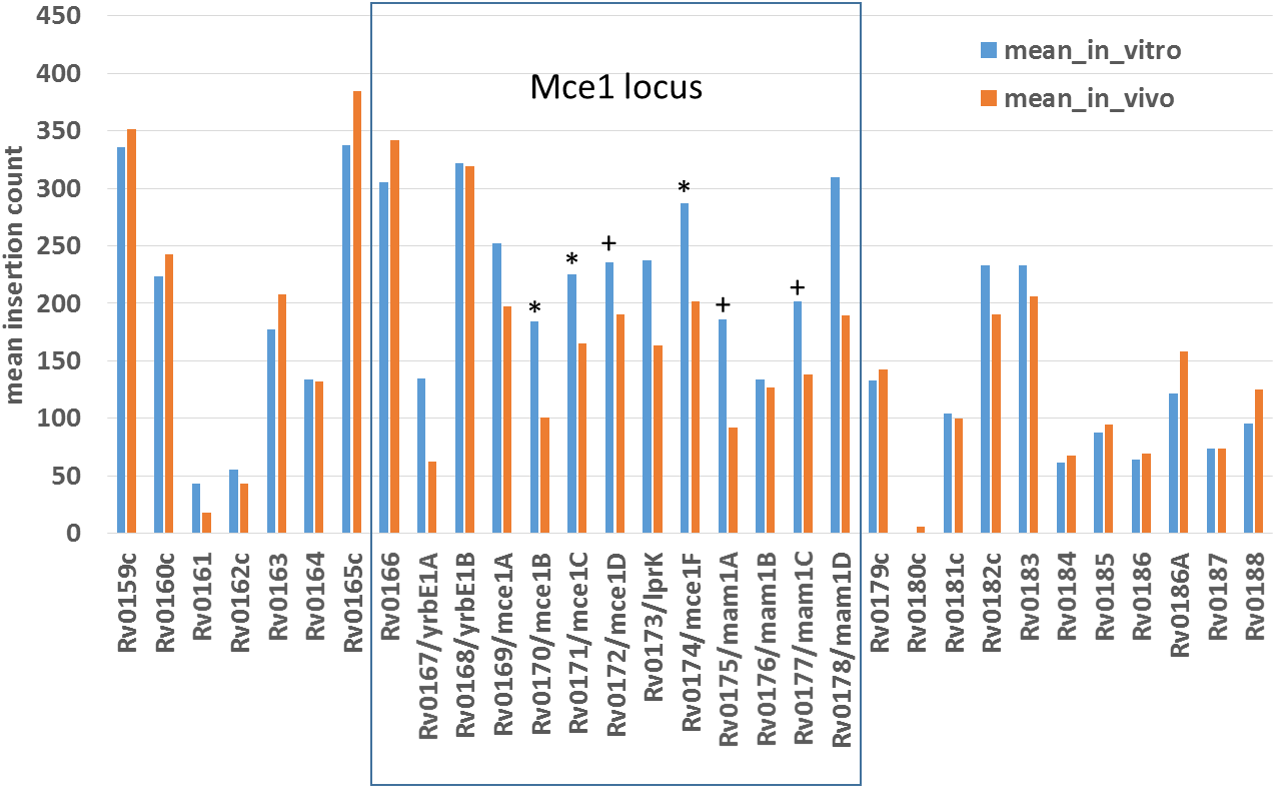
Reduction in mean insertion counts in-vivo (mice) for genes in the Mce1 locus. Genes that are detected as significant (q-value < 0.05) by ZINB regression are marked with ‘*’. Genes with marginal q-values of 0.05-0.11 are marked with ‘+’.

Similar examples include *esxB*, a secreted virulence factor, *fcoT* (thioesterase for non-ribosomal peptide synthase NRPS), *lysX* (lysinylation of cell-wall glycolipids [38]), *pitA* (involved in phosphate transport [39]), and *fadE33, hsaB* and *kshB,* which are involved in cholesterol catabolism [29]. All of these genes have been previously shown to be essential for infection in an animal model, but did not meet the threshold for significance based on resampling. The reason that several of these genes (like *fadE33* and *esxB*, shown in Figure 3) are detected by ZINB but not resampling is due primarily to changes in saturation; the non-zero mean (NZmean) changes only slightly, but the saturation drops significantly in each case; greater depletion of insertion mutants indicates reduced fitness. This highlights the value of treating the saturation parameter separately in the ZINB model. Another gene that exhibits this effect is SecA2. SecA2 is an alternative ATPase component of the *Sec* secretion pathway and is thought to help secrete other virulence factors inside the macophage [40]. SecA2 mutants have a weak phenotype in vitro (“growth defect” gene; [41]), so that the mean counts and saturation are low compared to other genes in-vitro (e.g. only 20% saturation, compared to ~50% globally); however, it becomes almost completely devoid of insertions in-vivo (Figure 3). While SecA2 was not detected as significant by either resampling or ANOVA, it was identified as conditionally essential by ZINB.

**Figure 3:**
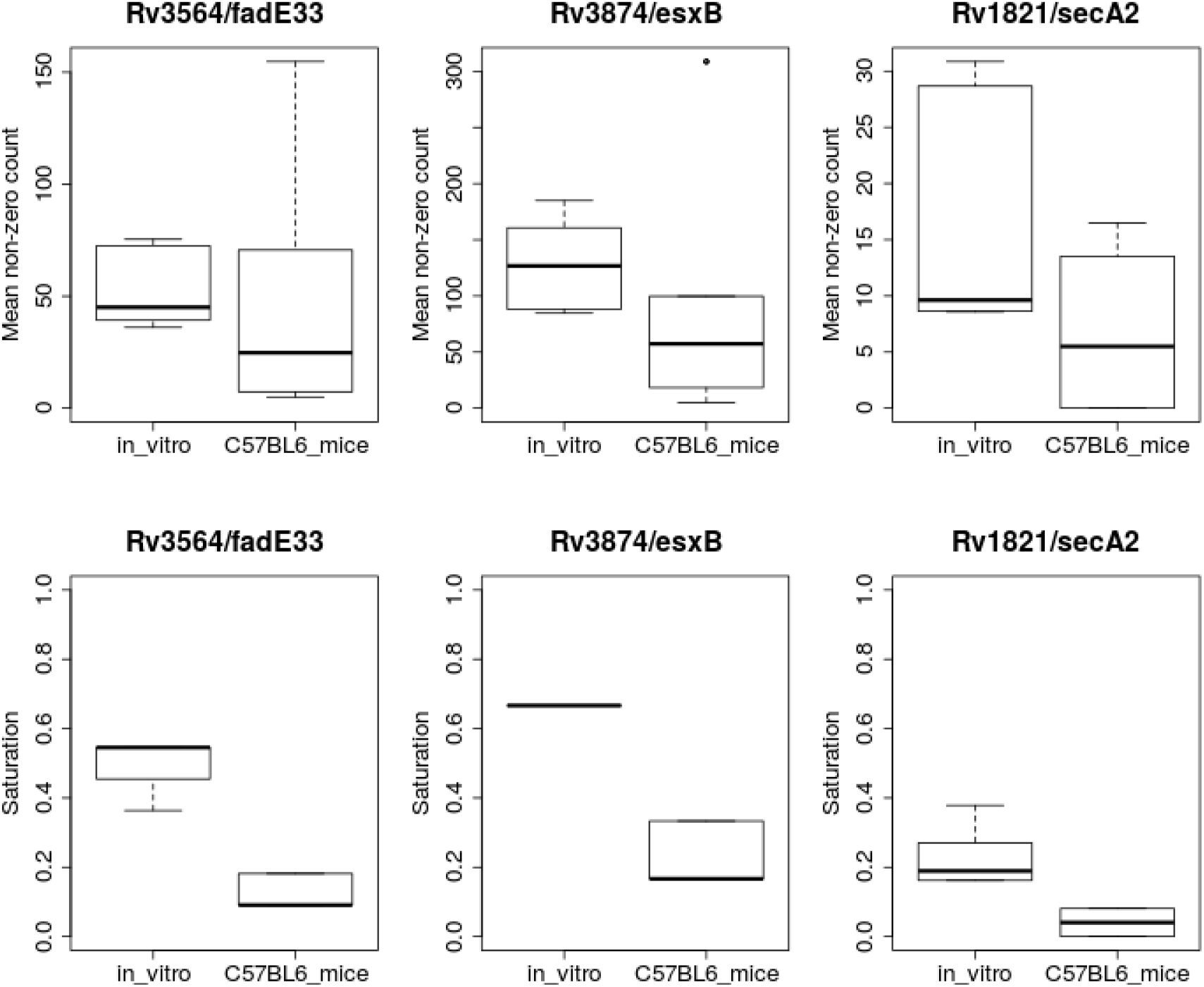
Statistics for three genes detected to vary significantly in mice compared to in-vitro based on ZINB regression, but not by resampling. The upper panels are the Non-Zero Mean (among insertion counts at TA sites with counts > 0), and the lower panels show the Saturation (percent of TA sites with counts > 0). Each box represents a distribution over 6 replicates.

Although ZINB identifies more genes (76) to be statistically significant than resampling on this dataset, it is unlikely that this excess is attributable to a large number of false positives. To evaluate the susceptibility of ZINB to generate false positives, we performed a comparison between replicates from the same condition by dividing the 6 in-vitro datasets into 2 groups (3+3). In this case, we expect to find no hits because there are (presumably) no biological differences. ZINB analysis identified only 15 genes as significantly different (*p_adj_* < 0.05), which suggests that the overall false positive rate for ZINB is quite low and probably reflects noise inherent in the data itself. Even resampling, when run on the same data (3 in-vitro vs. 3 in-vitro) for comparison, yielded 9 significant genes, which are presumably false positives.

### 3.3 Adjustment for Differences in Saturation Among Datasets

In real TnSeq experiments, it frequently happens that some datasets are less saturated than others. For example, there is often loss of diversity when passaging a Tn library through an animal model, possibly due to bottlenecking during infection or dissemination to target organs. TTR normalization was developed to reduce the sensitivity of the resampling method to differences in saturation levels of datasets. However, this type of normalization would be expected to exacerbate the detection of differences by ZINB. To compensate for this, we include offsets in the models that take into account the global level of saturation and non-zero mean for each dataset.

To evaluate the effect of the correction for saturation of datasets, we created artificially-depleted versions of some of the replicates analyzed in the previous Section (see Table 1). Specifically, for A1, A2, B1, and B2, we created “half-saturated” versions of each by randomly (and independently) setting 50% of the sites to 0. Since each of the original datasets had around 50% saturation to begin with, the half-saturated version have a saturation of approximately 25%.

Initially, we compared the original versions of A1 and A2 to B1 and B2 (scenario 1), with their observed level of saturation. The number of hits detected by ZINB (73) is similar to resampling (64). Recall that resampling with all 12 datasets yielded 186 significant genes; the number of hits is lower overall in this experiment because only 2 replicates of each were used, instead of 6. Then we compared fully-saturated versions of A1 and A2 to half-saturated B1 and B2 (scenario 2). ZINB-SA+ (with adjustment for saturation) identified nearly the same number of conditional essentials as resampling: 121 vs. 108. (see Table 2). The results are similar when half-saturated version of datasets A1 and A2 are used (scenario 3). However, when saturation adjustment is turned off, ZINB-SA^-^ produces dramatically more hits in case of wide saturation differences (2668 and 1139). The reason for this is that, by artificially reducing the saturation of either datasets A1 and A2 or B1 and B2, it amplifies the apparent differences in local saturation for many genes, to which ZINB is sensitive. The number of significant hits (conditional essentials) detected when half-saturated versions of all four datasets are used (scenario 4) is naturally lower (8 and 30), because there is much less information (fewer observations) available, making it more challenging for many genes to achieve statistical significance. Interestingly, when half-saturated versions of all four datasets are used, ZINB-SA^-^ works as expected, finding 37 hits (scenario 4), similar to resampling.

**Table 2:** Comparison of ZINB regression with and without saturation adjustment, for artificially-depleted samples.

| scenario | datasets compared | resampling | ZINB (SA<sup>+</sup>) | ZINB (SA<sup>-</sup>) |
| --- | --- | --- | --- | --- |
| 1 | (A1,A2) vs (B1,B2) | 64 | 73 | 162 |
| 2 | (A1,A2) vs (B1[50%],B2[50%]) | 108 | 121 | **2668** |
| 3 | (A1[50%],A2[50%]) vs (B1,B2) | 17 | 112 | **1139** |
| 4 | (A1[50%],A2[50%]) vs (B1[50%],B2[50%]) | 8 | 30 | 37 |

### 3.4 Application to Datasets with Multiple Conditions

In a prior study [21], a Himar1 transposon-insertion library in H37Rv was treated with sub-inhibitory concentrations of 5 different drugs: rifampicin (RIF), isoniazid (INH), ethambutol (EMB), meropenem (MERO), and vancomycin (VAN), all grown in 7H9 liquid medium. Combined with the untreated control, this makes 6 conditions, for which there were 3 replicate TnSeq datasets each (except INH; see Table 3). The TnSeq datasets had a high saturation of 60-65% (percent of TA sites with insertions). In the original analysis, each drug-treated sample was compared to the control using resampling [21]. Several conditionally essential genes were identified for each drug. Some genes were uniquely associated with certain drugs (for example, *blaC,* the beta-lactamase, was only required in the presence of meropenem), and other genes were shared hits (i.e. conditionally essential for more than one drug). Only one gene, *fecB,* was essential for all drugs, and its requirement for antibiotic stress tolerance was validated through phenotyping of a knock-out mutant.

**Table 3:** TnSeq datasets in different antibiotic treatments. MICs for H37Rv were obtained from [43].

| drug | concentration<br>($\mu\text{g/ml}$) | MIC<sub>50</sub><br>($\mu\text{g/ml}$) | # replicates | # of essentials<br>vs. untreated* |
| --- | --- | --- | --- | --- |
| untreated |  |  | 3 |  |
| isoniazid (INH) | 0.027 | 0.04 | 2 | 50 |
| rifampicin (RIF) | 0.004 | 0.02 | 3 | 68 |
| ethambutol (EMB) | 0.5 | 0.6 | 3 | 58 |
| meropenem (MER) | 1.2 | 30 | 3 | 106 |
| vancomycin (VAN) | 16 | 11 | 3 | 93 |

\* number of conditional essentials by comparison to the untreated condition using resampling

The raw datasets in this experiment have a number of sporadic outliers, consisting of isolated TA sites with observed insertion counts in one sample that are > 10 times higher than the others (even in other replicates of the same condition). Outliers can cause the appearance of artificial variability among conditions (inflating the mean count in one condition over the others in the ZINB model). Therefore, the raw datasets were normalized using the Beta-Geometric Correction (BGC) option in Transit, which is a non-linear transformation that reduces skew (extreme counts) in read-count distributions [42].

As a preliminary assessment, we did resampling of each drug condition against the untreated control, recapitulating the results in [21]. The number of conditional essentials is shown in Table 3. *fecB* was again observed to be the only hit in the intersection of all tests. We also observe other hits that can be rationalized, such as conditional essentiality of *blaC* (beta-lactamase) in presence of meropenem.

Next, variability among all 6 conditions was analyzed using several different methods. First, a simplistic but practical approach was taken by performing pairwise analyses of conditional essentiality using resampling (the permutation test for significant differences per gene in TRANSIT). For six conditions, there are 15 pairwise comparisons. Resampling was run independently on each pair of conditions, and the p-values were adjusted independently each time. By taking the union of conditionally-essential genes over all 15 pairwise comparisons, a total of 276 distinct genes was identified to have varying counts between at least one pair of conditions (Table 4).

**Table 4:** Identification of genes with significant variability across six conditions in antibiotic treatment data.

| method | # of varying genes |
| --- | --- |
| union of hits from 15 pairwise resampling tests | 276 |
| genes significant in any pairwise resampling<br>after pooled adjustment p-values | 267 |
| ANOVA | 234 |
| ZINB | 307 |

However, this straightforward approach is unfair because the p-values were adjusted independently. A more rigorous approach would be to perform resampling on all ~ 4000 genes for all 15 pairs of conditions, and then apply the p-value adjustment once on the pool of all ~ 60,000 p-values. When this is done, there are 267 significantly varying genes (using the lowest adjusted p-value for each gene). Thus, proper use of FDR correction results in a slightly more conservative list of hits.

The main problem with this approach is that it requires resampling to be run separately for all pairs of conditions, which does not scale up well as the number of conditions increases. As an alternative, ANOVA can be used to compare the counts across all six conditions simultaneously. When ANOVA is run (and the p-values are adjusted using the Benjamini-Hochberg procedure), only 234 significantly varying genes are identified. The 234 genes identified by ANOVA are almost completely contained in the set of those identified by pairwise resampling. Thus, ANOVA has lower sensitivity and under-reports genes with significant variability.

Finally, to identify genes that exhibit variability across all 6 conditions, we used ZINB regression. 307 genes were found to exhibit significant variation by ZINB, including genes identified in the original study, such as *fecB*, *blaC*, *pimE* (mannosyltransferase), and *secA2* (protein translocase) [21]. Another example of a gene found by both ZINB and pairwise resampling is *cinA* (Rv1901), which was specifically required for cultures exposed to sub-MIC concentrations of INH (Figure 5a). *cinA* is thought to be an NAD-dependent enzyme that plays a role in nucleoside recycling [44, 45], and thus it could confer tolerance to INH through a mechanism involving maintaining the intracellular NADH/NAD+ ratio [46].

**Figure 4:**
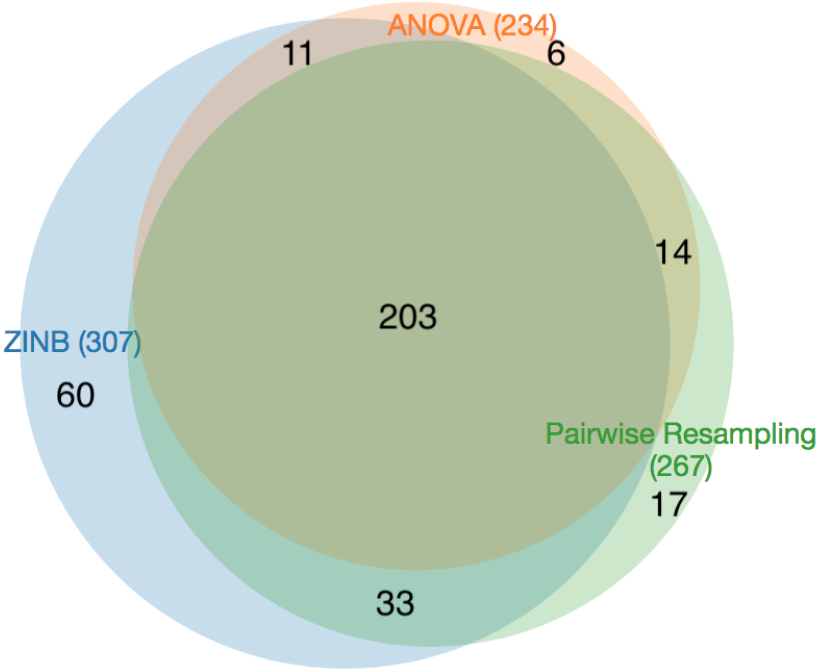
Venn diagram of genes with significant variability in different antibioitic treatments of transposon insertion counts evaluated by three different methods.

**Figure 5:**
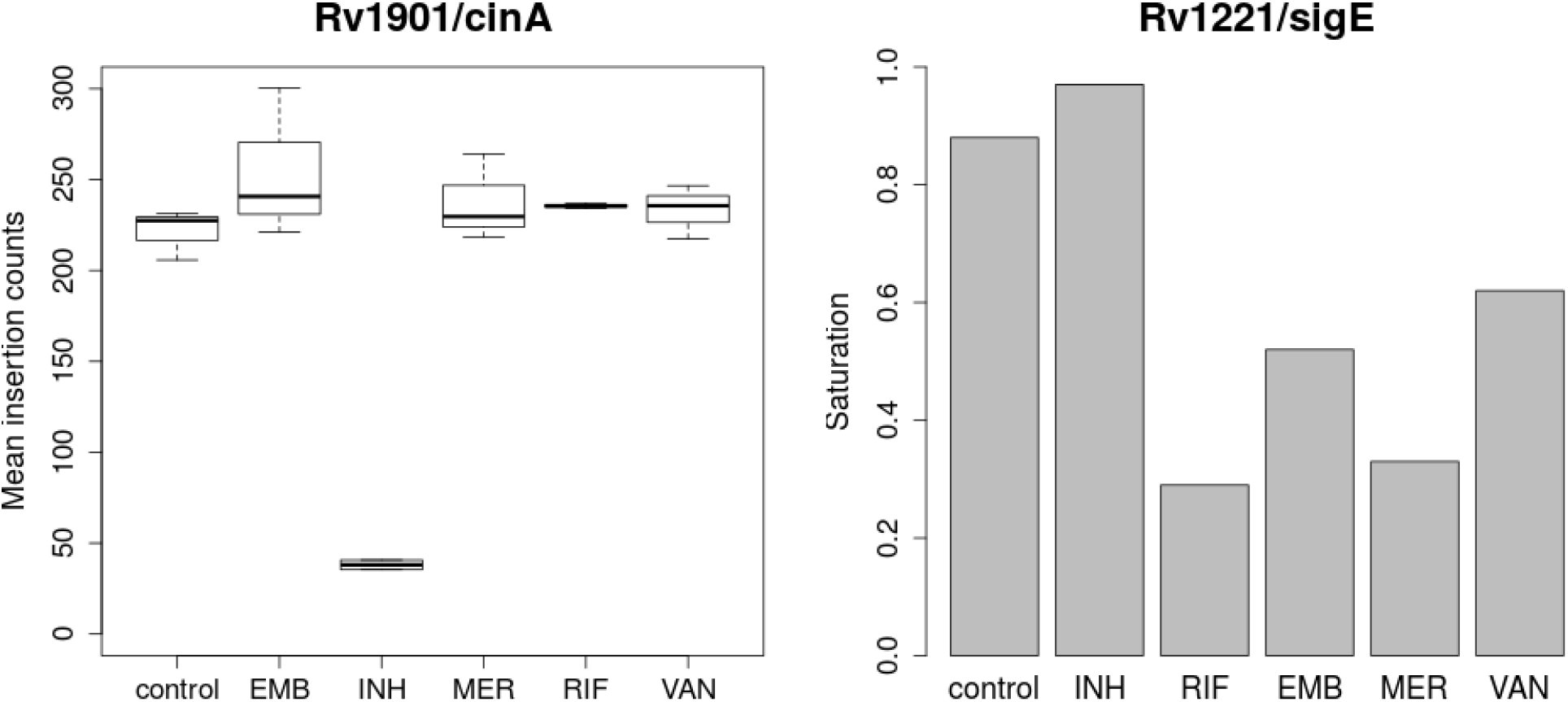
Significantly varying genes in cultures exposed to antibiotics. a) Mean insertion counts in CinA. b) Saturation in SigE (percent of TA sites with one or more insertions.

Compared to ANOVA, ZINB finds significantly more varying genes (307 compared to 234, 31% more). Put another way, ANOVA only identifies 76% of the genes with variability identified by ZINB. ZINB identified slightly more varying genes than pairwise resampling (71 additional genes). Many of these genes are on the margin and have adjusted p-value just slightly over the cutoff for resampling; 50% (36 out of 71 genes) have 0.05 < *p_adj_* < 0.2 for resampling. Among the remaining genes, one interesting case detected uniquely by ZINB is *sigE* (Figure 5b). While the mean insertion counts do not vary much for this gene (ranging between 17 and 27), the saturation level varies significantly among drug exposures, from nearly fully saturated in the control and INH conditions (88-97%), to highly depleted of insertions for RIF, MER and EMB (29 — 52%). This reduction suggests that *sigE* is required for tolerance of certain drugs. Indeed, this recapitulates the growth defects observed in a *ΔsigE* mutant when exposed to various drugs [47]. *sigE* is an alternative sigma factor that is thought to play a regulatory role in response to various stresses. This effect was only observable with a model that treats variations in saturation separately from magnitiudes of insertions.

## 4 Discussion

TnSeq has proven to be an effective tool for genome-wide assessment of functional requirements and genetic interactions in a wide range of prokaryotes. It is now being expanded to larger-scale experiments, such as profiling growth in media supplemented with an array of carbon sources or nutrients, or exposure to a variety of antibiotics/inhibitors, growth in a panel of different cell types, or infections in a collection of model-animals with different genetic backgrounds. Indeed, recent methods like BarSeq make such experiments efficient through barcoding of libraries, enabling highly multiplexed sequencing [48]. ZINB regression offers a convenient way of assessing variability of insertion counts across multiple conditions. It is more efficient than pairwise resampling (or permutation tests). Resampling is designed for two-way comparisons. Attempting to perform resampling between all pairs of conditions does not scale-up well, as the number of comparisons increases quadratically with number of conditions (for example, *n* = 20 conditions requires *n*(*n* – 1)/2 = 190 pairwise comparisons). In addition to the computational cost, there is a risk of loss of significance due to the p-value adjustment at the end, to control the overall false discovery rate. ZINB regression also performs better than ANOVA, a classic statistical test for conditional-dependence among observations from multiple groups. Our experimental results show that ANOVA is generally less sensitive than ZINB, detecting only a subset of varying genes, possibly because ANOVA relies on an assumption of normality [23]. Because most datasets are not fully saturated (due to lack of diversity of the library, bottlenecking, etc), TnSeq data usually has an over-abundance of zeros that cannot be approximated well with simpler distributions like Poisson or Binomial. The ZINB distribution, being a mixture model of a Negative Binomial and a zero component, allows the variance of the read-counts to be independent of the mean (unlike the Poisson) and allows sites with a count of zero to be treated separately (not all zeros are counted toward the mean). We showed with a likelihood ratio test that ZINB is a much more suitable model for TnSeq data (insertion counts) than ANOVA or NB (even when taking into account differences in the number of parameters).

To capture the conditional dependence of the parameters, the ZINB model is implemented as a regression model (with a log-link function), with vectors of coefficients to represent how the insertion counts vary across conditions. Thus the zero-component captures the changes in the level of saturation of a gene across conditions, and the NB component captures how the magnitudes of counts vary across conditions. Because of the zero-component included in the ZINB model, there is a risk that comparisons among datasets with different levels of saturation could result in a systematic inflation of the number of false positives (i.e. genes that look like they vary because of differences in the fraction of TA sites hit in different libraries). In fact, depending on the normalization procedure used, there can be a similar bias in the magnitudes of read counts that also causes more false positives when comparing datasets with widely-varying saturation. To compensate for this, we include “offsets” in the regression for the overall saturation and non-zero mean count for each dataset. Thus the coefficients learned in the model actually represent deviations in count magnitudes and saturation (local to each gene) relative to the genome-wide averages for each dataset. We showed in a synthetic experiment that failing to adjust for saturation differences leads to a large increase in the false-positive rate when comparing datasets with unbalanced levels of saturation. Moreover, when comparing replicates of the same condition against each other (which should not have any biological differences), we showed that ZINB detects almost no significantly varying genes, as expected, suggesting that it does not have a propensity to generate false positives. A potential limitation of ZINB, is that it can be sensitive to outliers. However, the impact of spurious high counts can be ameliorated by non-linear normalization methods like the Beta-Geometric correction [42], or other techniques like winsorization [49].

An important theoretical assumption made in the ZINB approach is that we model effects on the mean insertion counts at the gene-level, and treat differences among individual TA sites as random. Thus we pool counts at different TA sites within a gene, treating them as independent identically distributed (i.i.d.) samples. It is possible that different TA sites might have different propensities for insertion, for example, due to sequence-dependent biases. However, most Himar1 TnSeq studies to date have viewed the presence/abundance of insertions at TA sites as effectively random, resulting from stochastic processes during library construction (i.e. transfection), and no strong sequence biases have yet been identified. Early work on Himar1 transposon libraries in *E. coli* suggested that insertions were weakly influenced by local DNA bendability [50]. Subsequently, a small subset (< 9%) of TA sites in non-essential regions was found to be non-permissive for insertion, having the consensus (GC)GnTAnC(GC) [51]. But aside from these, no sequence bias has been found to explain differences in Himar1 insertions at different TA sites. In the future, if a sequence-dependent insertion bias were discovered, it is conceivable that the ZINB model could be modified to include conditional dependence on individual sites (or perhaps local sequence features). However, estimating counts at individual sites is subject to noise and likely to have high uncertainty, because, in many experiments, there are only one or two replicates of each condition, and hence only 1-2 observations per site. In the current approach, we pool counts from different TA sites in a gene when estimating the non-zero mean for each gene. An advantage of this simplification is that larger genes with more TA sites benefit from higher statistical confidence due to larger numbers of observations.

The significance of variability in each gene is determined by a likelihood ratio test, which identifies significantly variable genes based on the ability of using distinct parameters for each condition to increase the likelihood of the model, compared to a condition-independent null model (based on fitting parameters to the pooled counts, regardless of condition). A disadvantage of this approach is that the likelihood ratio test does not take into account certainty of the model parameter estimates. Therefore, Transit automatically filters out genes with insertions at only a single TA site (i.e. refuse to call them conditionally variable), because the coefficients of the model are too easily fit in a way that makes the likelihood look artificially high. By default our implementation requires at least 2 non-zero observations per condition to determine whether a gene exhibits significant variability across conditions. As with RNAseq, however, inclusion of multiple replicates increases the number of observations per gene, and this is a strongly recommended practice [25]. A more rigorous approach in Transit might be to apply a Wald test on the significance of the coefficients, which would also reveal cases where there are too few observations to be confident in the parameter estimates. More generally, a Bayesian approach might be better able to adjust (shrink) parameter estimates in cases of sparse data by combining them with prior distributions.

One advantage of the ZINB regression framework is that it can take into account additional information about samples in the form of covariates and interactions. This is commonly done in RNA-seq for experiments with more complex design matrices [52]. Examples include relationships among the conditions or treatments, such as class of drug, concentration, time of treatment/exposure, medium or nutrient supplementation, or genotype (for animal infections). By incorporating these in the model (with their own coefficients), it allows the model to factor out known (or anticipated) effects and focus on identifying genes with residual (or unexplained) variability. It can also be useful for eliminating nuisances like batch effects.

In theory, the ZINB regression method should work on TnSeq data from libraries generated with other transposons, such as Tn5 [1]. Tn5 insertions occur more-or-less randomly throughout the genome (like Himar1), but are not restricted to TA dinucleotides, though Tn5 appears to have a slight preference for insertions in A/T-rich regions [53]). Thus ZINB regression could be used to capture condition-dependent differences in magnitudes of counts or density of insertions in each gene. However, Tn5 datasets generally have much lower saturation (typically < 10%), since every coordinate in the genome is a potential insertion site, and thus the assumptions underlying the normalization procedure we use for Himar1 datasets (TTR) might not be satisfied for Tn5 datasets, requiring different normalization.

Of course, as with ANOVA, identifying genes that vary significantly across conditions is often just the first step and requires follow-up analyses to determine specific condition-dependent effects. For example, we observed that the NAD-dependent, nucleoside-recycling gene *cinA* was not just variable, but specifically required for tolerance of isoniazid. One could employ methods such as Tukey’s range test [54] to drill down and identify significantly different pairs of conditions. Another approach would be to use principle-component analysis (PCA) to uncover trends/patterns among TnSeq profiles and identify clusters of conditions producing similar effects genome-wide [55].

Our results establish the suitability of ZINB as a model for TnSeq data (insertion counts). Examples of genes where the phenotype is primarily observed in the saturation of the read-counts, such as SecA2 and SigE, highlight the advantage of modeling condition-dependent effects on both the magnitudes of counts in a gene and local level of saturation independently. Thus, ZINB regression is an effective tool for identifying genes whose insertion counts vary across multiple conditions in a statistically significant way.

## Conclusions

We have presented a novel statistical method for identifying genes with significant variability of insertion counts across multiple conditions based on Zero-Inflated Negative Binomial (ZINB) regression. The ZINB distribution was demonstrated to be appropriate for modeling transposon insertion counts because it captures differences in both the magnitudes of insertion counts (through a Negative Binomial) and the local saturation of each gene (through the proportion of TA sites with counts of 0). The method is implemented in the framework of a Generalized Linear Model, which allows multiple conditions to be compared simultaneously, and can incorporate additional covariates in the analysis. Thus it should make it a useful tool for screening for genes that exhibit significant variation of insertion counts (and hence essentiality) across multiple experimental conditions.

## Supporting information

Test Data and Example Files

## Abbreviations

TnSeq: transposon insertion mutant library sequencing
NB: Negative Binomial
ZINB: Zero-Inflated Negative Binomial
LRT: Likelihood Ratio Test
FDR: False Discovery Rate
TTR: Total Trimmed Read-count normalization
NZmean: Non-Zero mean
BGC: Beta-Geometric Correction
MIC: Minimum Inhibitory Concentration
CFU: Colony Forming Units

## Declarations

### Ethics approval and consent to participate

Treatment of mice used for experiementation in Section 3.2 was in accordance with the guidelines set forth by the Department of Animal Medicine of University of Massachusetts Medical School and Institutional Animal Care and Use Committee (IACUC; protocol #1649) and adhered to the laws of the United States and regulations of the Department of Agriculture. Mice were sacrificed by 5% isoflurane and cervical dislocation.

### Consent for publication

Not applicable.

### Availability of data and material

The methods described in this paper have been implemented in TRANSIT [15], which is publicly available on GitHub (http://saclab.tamu.edu/essentiality/transit/) and can be installed as a python package *(tnseq-transit*) using *pip.* The data from Section 3.2 (files with insertion counts from mouse infections), along with results files (spreadsheets with significant genes based on ZINB analysis), are provided in the Supplmental Material online.

### Competing interests

The authors declare that they have no competing interests.

### Funding

This work was supported by NIH grant U19 AI107774 (TRI, SE, DS, and CMS). The sponsor played no role in the design, interpretation, or writing of this manuscript.

### Authors’ contributions

SS implemented and tested the method described in this paper. MD and AS tested the method and editted the paper. CS performed the mouse infection experiment and reviewed the paper. RB, CS, SE, and DS provided critical input and feedback on the method and design of the experiments, and also editted the paper. TI developed, implemented, and tested the method, and wrote the paper. All authors read and approved the final manuscript.

## Acknowledgements

Not applicable.

## Figure Legends

### Additional Files

- **Supplemental Table 1** - ZINB output file (spreadsheet, tab-separated format) containing results from Section 3.2 on comparison of a TnSeq library for *M. tuberculosis* H37Rv grown in-vitro versus in C57BL/6 mice. For each ORF in the genome, an analysis of the mean, NZ-mean, and saturation in each condition is given, along with p-value and adjusted p-value. Significantly varying genes are those with *p_adj_* < 0.05.
- **Supplemental Table 2** - ZINB output file containing results for Section 3.4 on comparison of a TnSeq library for *M. tuberculosis* H37Rv grown in presence of five antibiotics, isoniazid (INH), rifampicin (RIF), ethambutol (EMB), meropenem (MER), and vancomycin (VAN).
- **Data from Mouse Infections, and ZINB Test Example (zip file)** - Experimental data for the mouse experiment in Section 3.2 (.wig files with transposon insertion counts at TA sites), along with Instructions (README.docx) on how to do the ZINB analysis in TRANSIT (commands and output files).

## Notes

#### Summary of Updates

Improved the notation and formalism in the Methods section. Extensively re-wrote the Discussion to focus more on strengths and limitations.

