## Supplementary material for "Statistical Analysis of Variability in TnSeq Data Across Conditions Using Zero-Inflated Negative Binomial Regression": Test Data and Example Files: README.docx

ZINB Test Example

In this example, ZINB regression in TRANSIT will be used to compare 6 replicates of a TnSeq library of *M. tuberculosis* H37Rv grown in vitro to 6 replicates of the same library infected into C57BL/6 mice. The input files consist of 'wig files' (A1.wig-A6.wig, and B1.wig-B6.wig), which contain counts of insertions at TA sites in genome. The reference genome is H37RvBD1.fna (fasta), and the accompanying annotation file defining ORFs is H37RvBD1.prot_table.

Step 1. Download and install TRANSIT.

If you are using Linux, a simple version of the instructions is to run 'pip install tnseq-transit'. Note that this will install a number of dependencies for you (if not already installed), such as Numpy, wxPython, etc. You also have to have R installed on your machine. Alternatively, TRANSIT source code can be downloaded and installed from GitHub. For more thorough details on installing TRANSIT, please see the Installation instructions on: https://transit.readthedocs.io/en/latest/transit_install.html

Step 2. Create a combined wig file. Open up a shell or terminal and type the following command-line:

> transit export combined_wig A1.wig,A2.wig,A3.wig,A4.wig,A5.wig,A6.wig,B1.wig,B2.wig,B3.wig,B4.wig,B5.wig,B6.wig H37RvBD1.prot_table IV_combined_wig_TTR.txt

Note that all 12 wig filenames are given as a single comma-separated string. This will create a tab-separated output file (IV_combined_wig_TTR.txt) with insertion counts at TA sites for each dataset. (Coords of TA sites are given in column 1). In the process, each dataset is normalized by TTR.

(Depending on how it was installed, you might have to type 'transit.py', or perhaps 'python <PATH_TO_TRANSIT_DIR>/src/transit.py'.)

Step 3. Run the ZINB analysis. Open up a shell or terminal and type the following command-line:

> python transit zinb IV_combined_wig_TTR.txt samples_metadata.txt H37RvBD1.prot_table IV_ ZINB_IV.txt

The output file (ZINB_IV.txt), with statistical analysis for each gene, is in tab-separated format and can be opened as a spreadsheet. The columns are:

| Rv | ORF id |
| --- | --- |
| Gene | gene name, if defined |
| TAs | number of TA sites in the gene (sites in the N- and C-terminal 5% of the ORF are trimmed by default) |
| mean_in_vitro | mean insertion count for gene averaged over TA sites and replicates for A1-A6 |
| mean_in_vivo | mean insertion count for gene averaged over TA sites and replicates for B1-B6 |
| NZmean_in_vitro | mean insertion count for averaged over non-zero sites for A1-A6 |
| NZmean_in_vivo | mean insertion count for averaged over non-zero sites for B1-B6 |
| NZperc_in_vitro | fraction of sites in gene with non-zero values for A1-A6 |
| NZperc_in_vivo | fraction of sites in gene with non-zero values for B1-B6 |
| pval | p-value from ZINB analysis |
| padj | adjusted p-value by Benjamini-Hochberg FDR-correction |
| status | flag indicating reason for genes for which analysis was not applied |

Genes with significant variability between in-vitro (A) and in-vivo (B) conditions are defined as those with padj<0.05. You might want to sort the rows by the 'padj' column to rank the hits at the top.
